# Sequence and structural variations determining the recruitment of WNK kinases to the KLHL3 E3 ligase

**DOI:** 10.1101/2020.06.29.178285

**Authors:** Zhuoyao Chen, Jinwei Zhang, Raphael Heilig, Fiona J. Sorrell, Vincenzo D’Angiolella, Roman Fischer, Dario R. Alessi, Alex N. Bullock

## Abstract

The BTB-Kelch protein KLHL3 is a Cullin3-dependent E3 ligase that mediates the ubiquitin-dependent degradation of kinases WNK1-4 to control blood pressure and cell volume. A crystal structure of KLHL3 has defined its binding to an acidic degron motif containing a PXXP sequence that is strictly conserved in WNK1, WNK2 and WNK4. Mutations in the second proline abrograte the interaction causing the hypertension syndrome pseudohypoaldosteronism type II. WNK3 shows a diverged degron motif containing 4 amino acid substitutions that remove the PXXP motif raising questions as to the mechanism of its binding. To understand this atypical interaction, we determined the crystal structure of the KLHL3 Kelch domain in complex with a WNK3 peptide. The electron density enabled the complete 11-mer WNK-family degron motif to be traced for the first time revealing several conserved features not captured in previous work, including additional salt bridge and hydrogen bond interactions. Overall, the WNK3 peptide adopted a conserved binding pose except for a subtle shift to accommodate bulkier amino acid substitutions at the binding interface. At the centre, the second proline was substituted by WNK3 Thr541, providing a unique phosphorylatable residue among the WNK-family degrons. Fluorescence polarisation and structural modelling experiments revealed that its phosphorylation would abrogate the KLHL3 interaction similarly to hypertension-causing mutations. Together, these data reveal how the KLHL3 Kelch domain can accommodate the binding of multiple WNK isoforms and highlight a potential regulatory mechanism for the recruitment of WNK3.

## Introduction

WNK (with no lysine) kinases play a key role in mammalian blood pressure regulation. WNKs, together with their downstream targets SPAK (SPS1-related proline/alanine-rich kinase) and OSR1 (oxidative stress-responsive kinase 1), regulate cation-chloride channels to achieve ion homeostasis in the kidney and neurons through a cascade of phosphorylation [1]. Four WNK isoforms (WNK1-4) are activated by autophosphorylation on a conserved serine in the kinase domain activation segment (WNK1 Ser382) upon exposure of cells to hypotonic and low [Cl^−^] conditions [2]. Activated WNK kinases then stimulate the kinase activity of SPAK/OSR1 by phosphorylating a conserved threonine residue in their kinase activation segments (SPAK Thr233, OSR1 Thr185) [3], which is facilitated by interaction between the SPAK/OSR1 CCT (Conserved C-terminal) domain and WNK RFXV/I peptide motifs [2]. Similarly, the SPAK/OSR1 CCT domain is recruited to RFXV/I motifs in the N-terminus of N[K]CC cation-chloride channels, and thereby phosphorylates conserved threonine residues in their cytoplasmic domains to activate channel activity [4]. Conversely, phosphorylation on KCC cation-chloride channels by the WNK-SPAK/OSR1 axis has an inhibitory function [5]. Excessive activity of the WNK-SPAK/OSR1 cascade is the primary cause of PHAII (pseudohypoaldosteronism type II) hypertension and hyperkalemia syndrome [6].

Further regulation of WNK activity and blood pressure is provided by the KLHL3 E3 ligase, which degrades WNK1-4 through the ubiquitin-proteosome system [7, 8]. KLHL3 is a BTB-Kelch family protein that forms the substrate adaptor of a Cullin3-dependent RING E3 ligase (CRL3) complex also consisting of Cullin3, RBX1 and NEDD8. As a substrate adaptor, KLHL3 recruits WNK1-4 through its C-terminal Kelch domain, whereas the N-terminal BTB domain binds to Cullin3 [9, 10]. The RING domain protein RBX1 also binds to Cullin3 and serves to recruit E2-ubiquitin conjugates to the CRL3 complex before transfer of the ubiquitin onto WNK substrates. This substrate ubiquitination is catalysed by neddylation of Cullin3, which induces a conformation favourable for conjugation [11, 12]. Mutations in KLHL3 that disrupt interactions with either the WNK kinases or Cullin3 lead to WNK upregulation and the development of PHAII hypertension and hyperkalemia syndrome [7, 8, 13, 14]. Similarly, mutations within a WNK4 non-catalytic motif (residues 557-567) also cause PHAII hypertension syndrome [4, 6] and have been found to occur within an acidic degron motif for recognition by KLHL3 [15]. The structural basis for the interaction of KLHL3 and WNK4 has been recently determined and clearly defines the disruptive effects of PHAII-associated mutations in the complex interface [9].

The acidic degron motif in WNK4 is conserved among WNK1-4 allowing all four WNK family proteins to bind to KLHL3 [9]. However, WNK3 is notably distinguished by the presence of four amino acid substitutions in its acidic degron motif compared to the 11-mer WNK4 degron that was previously crystallized in complex with KLHL3 (Figure 1). These include substitutions at two proline sites in WNK4 that have potential to alter the peptide backbone. In addition, the WNK3 degron uniquely contains a threonine residue that potentially allows for a phosphorylation-dependant regulatory mechanism.

**Figure 1.**
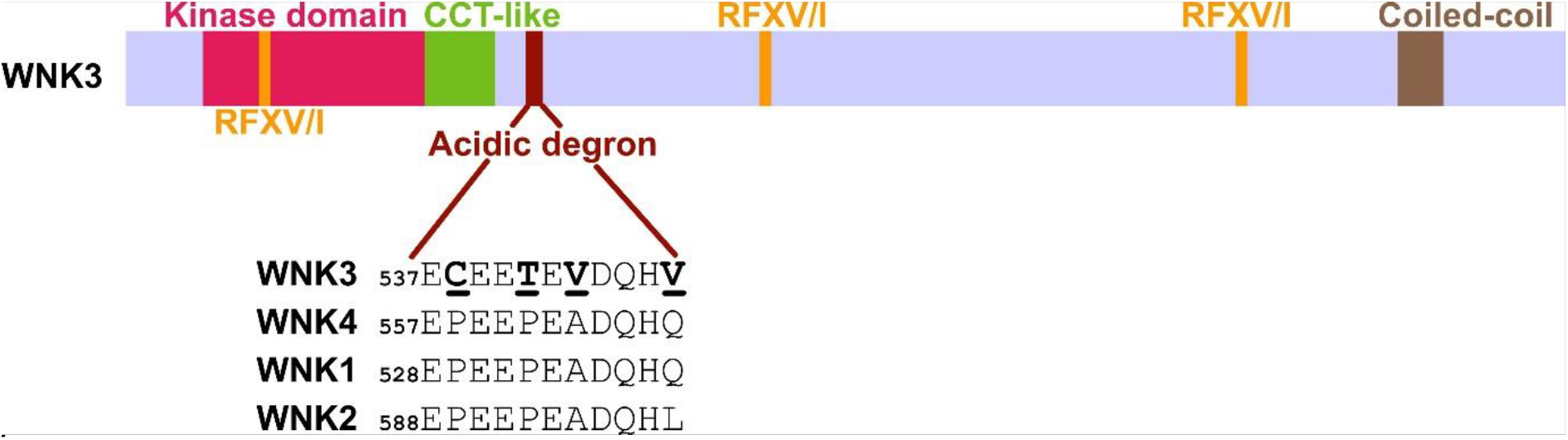
Schematic diagram showing the typical domain composition of WNK family kinases. In addition to an N-terminal kinase domain, WNK1-4 contain an acidic degron motif that mediates their recruitment to the KLHL3 E3 ligase. Degron sequences are shown for each family member. The WNK3 degron motif contains four amino acid substitutions highlighted with bold underlined letters. Coil indicates a C-terminal coiled-coil domain. CCT-like is a circular permutant of the CCT domain of the OSR1 and SPAK kinases. WNK kinases contain additional RFXV or RFXI motifs (where X denotes any amino acid) that mediate their recruitment to the CCT domain of SPAK/OSR1.

To address these differences, we report the 2.8 Å structure of the KLHL3 Kelch domain in complex with the WNK3 degron motif, as well as peptide binding assays that characterise the interactions of KLHL3 with variant WNK3 peptides. The new structure reveals a conserved structural mechanism for WNK family interaction with KLHL3 that explains the interactions of the four amino acid differences in WNK3. The structure and peptide binding assays also reveal that threonine phosphorylation in the WNK3 degron would disrupt the complex interface providing a potential regulatory mechanism for the KLHL3-WNK3 interaction.

## Materials and Methods

### Recombinant protein expression and purification

For binding assays, the kelch domain of human KLHL3 (a.a. 290-587) was cloned into the bacterial expression vector pGEX6P-1 (MRC-PPU plasmid DU44387) and expressed as a GST fusion protein as previously described [9]. Briefly, *E. coli* BL21 cells were transformed with plasmid DNA and grown at 37°C in LB broth containing 100 μg/mL ampicillin until OD_600_ reached 0.6. Cells were then cultured at 16°C for a further 16 h in the presence of 50 μM IPTG for protein induction. Harvested cells were lysed by sonication and the lysate clarified by centrifugation. GST fusion protein was affinity purified using 0.5 mL glutathione–sepharose beads and eluted in buffer (50 mM Tris-HCl pH 7.5, 150 mM NaCl, 2 mM DTT) containing 20 mM glutathione.

For structure determination, human KLHL3 (a.a. 298–587) was cloned into the pNIC28-Bsa4 vector (Addgene plasmid #110251), which provides an N-terminal hexahistidine tag as previously described [9]. Expression was performed in BL21(DE3)-pRARE cells grown in LB broth containing 50 μg/mL kanamycin and 50 μg/mL chloramphenicol. When OD_600_ reached 0.6, expression was induced overnight at 18°C by addition of 0.4 mM IPTG. Cell pellets were lysed by sonication and recombinant protein purified using nickel-sepharose beads equilibrated in binding buffer (50 mM HEPES pH 7.5, 500 mM NaCl, 5% glycerol, 10 mM imidazole, 1 mM TCEP). Protein was eluted stepwise in buffers containing 100-500 mM imidazole. Further purification was achieved by size exclusion chromatography on a HiLoad 16/600 Superdex 200 pg column. The hexahistidine tag was cleaved using TEV protease and the correct mass of KLHL3 was confirmed by LC-MS mass spectrometry. Protein was stored in 50 mM HEPES pH 7.5, 300 mM NaCl, 0.5 mM TCEP, 2 mM DTT.

### Fluorescence polarisation

Fluorescence polarisation measurements were performed at 25°C using KLHL3 kelch domain (GST-KLHL3, a.a. 298–587(end) (DU44387)) buffered in 50 mM Tris-HCl pH 7.5, 150 mM NaCl, 2 mM DTT. All peptides (ECEETEVDQHV, WNK3 a.a. 537–547) [EP5603], (ECEET(P)EVDQHV, WNK3 a.a. 537–547, Thr541 is phosphorylated) [5604], and (ECEEEEVDQHV, WNK3 a.a. 537–547, Thr541 is mutated to Glu) [EP5605] were conjugated to the Lumio green fluorophore via an N-terminal linker (CCPGCCGGGG) and dialysed into assay buffer before use. Samples were prepared with 10 nM Lumio-green labelled peptide and the indicated concentration of protein in a final volume of 30 μL. Fluorescence polarisation measurements were recorded using a BMG PheraStar plate reader, with an excitation wavelength of 485 nm and an emission wavelength of 538 nm, and measurements were corrected to the fluorescent probe alone. Data analysis and graphing were performed in GraphPad Prism6. One Site Specific binding with Hill Slope was assumed (model Y=B_max_*X^h/*K*_d_^h + X^h) and the disassociation constant, and associated standard error was obtained. All experimental bindings were repeated at least twice and comparable results to those shown in the present study were obtained.

### Crystallisation and structure determination

KLHL3 was concentrated to 9 mg/mL using a 10 kDa centrifugal concentrator (Millipore) and mixed with 2 mM WNK3 peptide (ECEETEVDQHV, synthesised by Severn Biotech Ltd). Crystallisation was performed using sitting drop vapour diffusion. Initial crystals obtained from coarse screens were used to make seed stocks for further fine screening. The best-diffracting crystals of the KLHL3-WNK3 complex were obtained at 4°C by mixing 20 nL seed stock with 75 nL of protein and 75 nL of a reservoir solution containing 6% PEG4K, 0.1 M acetate pH 5.1. Prior to vitrification in liquid nitrogen, crystals were cryoprotected by direct addition of reservoir solution supplemented with 25 % ethylene glycol. Diffraction data were collected on beamline I03 at the Diamond Light Source, Didcot, UK. Data were processed in PHENIX [16]. Molecular replacement was performed with Phaser MR in Phenix using PDB code 4CH9 chain A (kelch domain of KLHL3) as the model. COOT [17] was used for manual model building and refinement, whereas PHENIX.REFINE [18] was used for automated refinement. TLS parameters were included at later stages of refinement. Structure factors and co-ordinates have been deposited in the PDB with accession code 5NKP.

### Tissue culture and phosphorylation site mapping by LC-MSMS

HEK293T cells were cultured in Dulbecco’s modified Eagle’s medium (DMEM, Gibco/Invitrogen) supplemented with 10% fetal bovine serum (FBS, Sigma Aldrich), 100 U/mL penicillin sodium and 100 lg/mL streptomycin sulphate (Sigma Aldrich) in a humidified incubator at 37°C with 5% CO_2_. For phospho-mapping experiments, full length human WNK3 (isoform 2) was cloned into a pCMV5-FLAG vector providing an N-terminal Flag affinity tag (MRC-PPU plasmid DU4949). Cells were split into 150 mm dishes and transfected with 14 μg Flag-WNK3 construct using polyethylenimine (Polysciences). Some 36 hours post-transfection, culture medium was removed and the cells were treated with either isotonic buffer or hypotonic buffer to allow for phosphorylation under different stimuli. Isotonic high potassium buffer contained 20 mM HEPES pH 7.4, 95 mM NaCl, 50 mM KCl, 1 mM CaCl_2_, 1 mM MgCl_2_, 1 mM Na_2_HPO_4_, 1 mM Na_2_SO_4_. Hypotonic high potassium buffer contained 20 mM HEPES pH 7.4, 80 mM KCl, 1 mM CaCl_2_, 1 mM MgCl_2_, 1 mM Na_2_HPO_4_, 1 mM Na_2_SO_4_. Following 30 min incubation, cells were washed in PBS buffer and harvested for analysis. WNK3 was immunoprecipitated by anti-Flag M2 affinity resin (Sigma-Aldrich, A2220) and eluted with 3xFlag peptide. Protein samples were digested in elastase after DTT reduction, iodoacetamide alkylation and methanol-chloroform precipitation. The samples were then desalted using a SOLA HRP SPE Cartridge (Thermo Fisher, 60109-001) following the manufacturer’s instruction.

Samples were analysed on a LC-MS/MS platform consisting of a Dionex Ultimate 3000 UPLC and Orbitrap Fusion Lumos mass spectrometer (both Thermo Fisher). Peptides were separated in a 60 min gradient from 2% ACN/5% DMSO in 0.1% Formic Acid to 35% CAN in the same buffer on a 50 cm Easy-Spray Column (Thermo Fisher). MS spectra were acquired in the ion trap with a resolution of 120,000 and with an ion target of 4e5. MS/MS spectra were acquired in the ion trap in rapid mode after HCD fragmentation. Maximum injection time was set to 35 ms and AGC target was 4000. Selected precursors were excluded for 60 seconds.

LC-MS/MS data were analysed with PEAKS 7.0 against the Uniprot database using an Elastase digest pattern and 10 ppm/0.5 Da (MS/ MS/MS) mass tolerance. Peptide level false discovery rate was adjusted to 1% and modifications were set according to the experimental parameters described above.

## RESULTS

### The WNK3 acidic degron motif binds potently to KLHL3 but does not bind if phosphorylated at Thr541

To investigate the binding of WNK3 variants to the KLHL3 kelch domain we employed a fluorescence polarisation assay developed previously to map the degron motif of WNK4 (^557-^EPEEPEADQHQ) [9]. In this assay, peptides containing the WNK family degron motif were conjugated to a Lumio green fluorophore and titrated with recombinant KLHL3 kelch domain. Previous measurements indicated that the paralogs WNK1-4 could bind to KLHL3 with similarly high affinity (*K*_D_ values of 0.3 – 0.9 μM) [9]. In agreement, we observed here that the 11-mer WNK3 degron motif (ECEETEVDQHV) bound to KLHL3 with *K*_D_ = 0.67 μM (Figure 2). To examine whether modifications of the central Thr541 residue could form a potential regulatory mechanism we prepared an equivalent WNK3 peptide carrying a single phosphorylation at Thr541. Strikingly, the binding of this phospho-WNK3 peptide was too severely destabilized to measure a dissociation constant (*K*_D_ > 50 μM, Figure 2). A similar loss of KLHL3 interaction was observed using a WNK3 T541E mutant degron (Figure 2). Together, these results indicated that WNK3 could bind to KLHL3 with similar potency to WNK4, and that WNK3 binding was abrogated upon modification of WNK3 Thr541.

**Figure 2.**
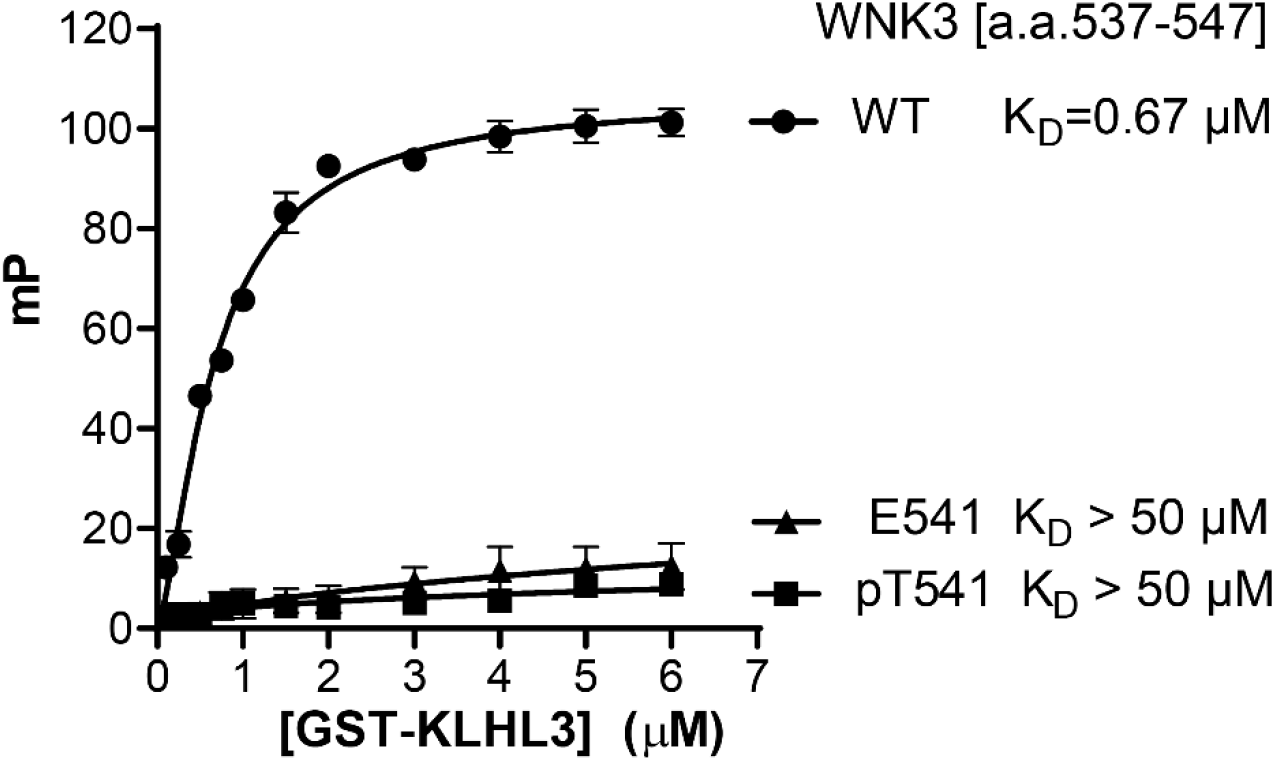
Analysis of the interaction between KLHL3 and WNK3 variants by fluorescence polarisation. Purified GST-KLHL3 a.a. 298–587(end) was diluted appropriately and mixed at a 1:1 volume ratio with 20 nM Lumino-Green-labelled WNK3 peptides to the concentration stated in the Figure, with the peptide concentration consistent at 10 nM. Fluorescence polarisation measurements were recorded and corrected to the fluorescent probe alone. Each data point represents three technical replicates. One site-specific binding with hill slope was assumed and the disassociation constant was obtained. Binding curves, assuming one-site-specific binding, were then generated with Prism6 using milli-polarization (mP) units.

### Structure determination of the WNK3 acidic degron motif in complex with KLHL3

To investigate whether the divergent sequence of the WNK3 degron impacts upon its binding mode, we determined a 2.8 Å crystal structure of the 11-mer WNK3 degron in complex with the KLHL3 kelch domain (Figure 3). Data processing and refinement statistics are presented in Table 1. The co-structure was solved in space group C2 2 2_1_, with two WNK3 complexes in the asymmetric unit. The entire WNK3 peptide was traced in chain C (Figure 4A), but electron density was not resolved for the C-terminal residue Val547 in chain D (Figure 4B). In both WNK3 chains, Cys538 was observed to form an intermolecular disulphide-bond that potentially contributed to the observed crystal lattice. Despite distinct crystal contacts, the KLHL3 structure and WNK3 binding mode were highly conserved with the KLHL3-WNK4 complex (PDB 4CH9) suggesting a true representation of the physiologically-relevant assembly.

**Table 1.** Data collection and refinement statistics. Values in parentheses indicate data for the highest resolution shell.

| Parameter | KLHL3-WNK3 peptide |
| --- | --- |
| **Data collection** |  |
| Space group | C 2 2 21 |
| Cell constants |  |
| a, b, c (Å) | 84.66, 169.72, 123.86 |
| $\alpha$, $\beta$, $\gamma$ (°) | 90, 90, 90 |
| Resolution (Å) | 50.02 - 2.8 (2.9 - 2.8) |
| Unique observations | 22366 (2197) |
| Completeness (%) | 99.98 (100.00) |
| R<sub>merge</sub> | 0.09155 (0.4966) |
| I/$\sigma$ | 8.18 (1.51) |
| **Refinement** |  |
| Resolution (Å) | 50.02 - 2.8 |
| MR model | 4CH9 |
| Copies in ASU | 2 |
| R<sub>work</sub>/R<sub>free</sub> | 0.2274/0.2496 |
| Number of atoms | 4566 |
| Average B-factor (Å<sup>2</sup>) | 38.1 |
| Average B-factor (protein) (Å<sup>2</sup>) | 38.1 |
| Average B-factor (peptide) (Å<sup>2</sup>) | 43 |
| Average B-factor (solvent) (Å<sup>2</sup>) | 37.5 |
| RMSD (bonds) (Å) | 0.007 |
| RMSD (bonds) (°) | 0.75 |

**Figure 3.**
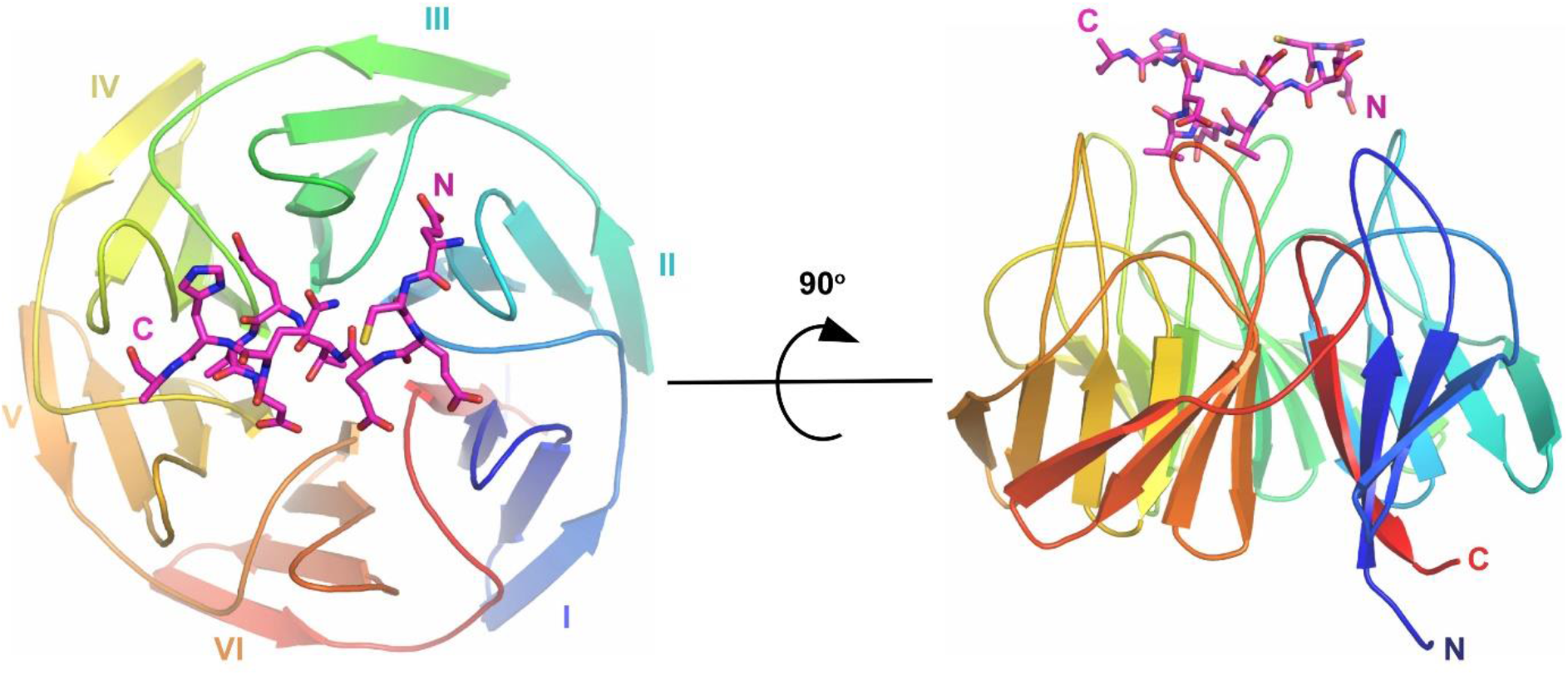
Crystal structure of the KLHL3 Kelch domain bound to WNK3 peptide determined at 2.8 Å. Overview of the structure of the KLHL3 Kelch domain (rainbow ribbon) in complex with WNK3 peptide (purple sticks). Kelch repeats forming blades I to VI are labelled. N and C termini for both KLHL3 and WNK3 are labelled.

**Figure 4.**
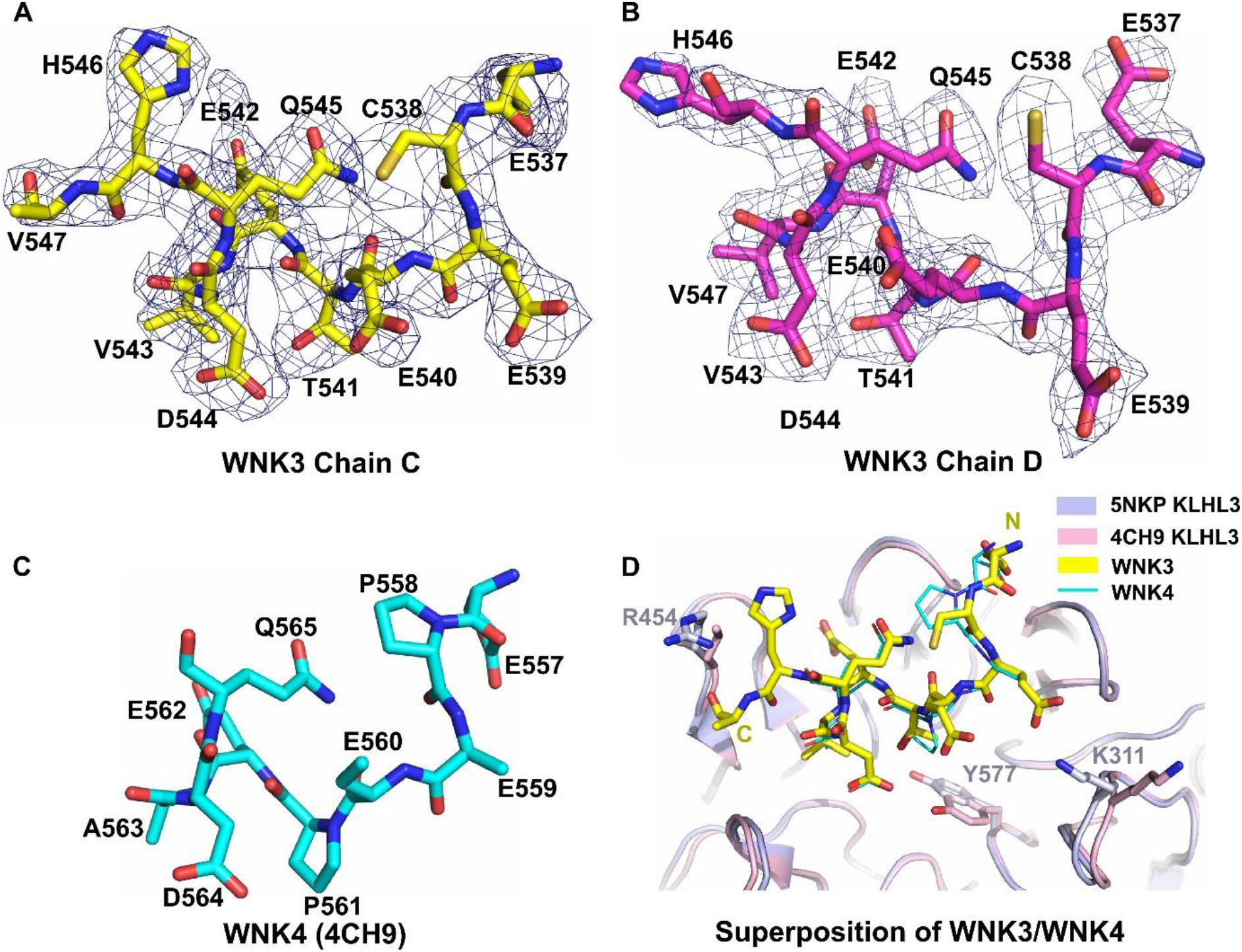
Structural comparison of WNK family degrons. Stick representation and 2Fo-Fc electron density maps contoured at 1.0 σ for (**A**) WNK3 chain C and (**B**) WNK3 chain D. WNK3 residues and their numbers are labelled (PDB 5NKP). (**C**) Stick representation of WNK4 degron peptide from PDB 4CH9. Residues are labelled as shown. (**D**) Superposition of the KLHL3-WNK3 and KLHL4-WNK4 complexes (PDBs 5NKP and 4CH9, respectively). The Kelch domains of KLHL3 are illustrated in ribbon representation. KLHL3 residues that adopt different conformations are highlighted with stick representation.

Consistent with other family structures, the KLHL3 kelch domain consists of six propeller blades arranged along a central axis. Each blade is folded as a twisted β-sheet comprising four anti-parallel beta-strands. The WNK3 peptide sits atop the substrate binding pocket, which is shaped by the six loops connecting the β2-β3 strands, as well as the extended β4-β1 loops that connect adjacent blades. Near identical conformations are observed for the peptide backbone in the two WNK3 chains, as well as side chains in the core interface, but several solvent-exposed side chains appear free to adopt alternative conformations, including Glu537, Cys538, Glu540 and His546 (Figures 4A and 4B).

### Structural comparison of WNK3 and WNK4 binding to KLHL3

Of note for structural comparisons, the WNK3 and previous WNK4 co-structures were both determined under acidic conditions (pH 5.1 for WNK3 and pH 4.3 for WNK4), providing similar environments for electrostatic interactions. Overall, the bound peptides adopt a conserved conformation, containing an extended N-terminal segment that spans the two proline positions in WNK4, and a C-terminal segment that folds into a single helical turn (Figures 3 and 4). The conserved glutamine in the WNK family degron motif acts to stabilize this turn through an intramolecular hydrogen bond, which in WNK3 is mediated between the side chain of Gln545 and the backbone carbonyl of Glu540 (Figures 4B and 4C).

Key differences in the two structures include the substitutions of WNK3 Cys538 and Thr541 at the two proline positions of WNK4. While the proline phi-psi angles are near ideal for extended structure (near −60° and +135°, respectively), the substitutions in WNK3 are associated with a subtle shift (up to 1 Å) in the peptide backbone position. This shift appears favourable for the binding interface to accommodate the bulkier WNK3 Val543 substitution for WNK4 A563, as well as the branched Thr541 side chain for WNK4 Pro561.

A number of polar and hydrophobic interactions in the KLHL3-WNK3 structure are strongly conserved with the equivalent WNK4 complex. Perhaps the most important interaction is the salt bridge formed between WNK3 Asp544 and KLHL3 Arg528 (Figure 5). The substitution R528H in KLHL3 is the most frequent mutation in Gordon’s hypertension syndrome and is sufficient to abolish KLHL3 function [8, 9]. WNK3 Val543 is a conservative change from WNK4 Ala563 and packs adjacent to this salt bridge to form equivalent van der Waals interactions with KLHL3 Y449 and H498. The preceding WNK3 residue Glu542 is also buried at the KLHL3 surface between KLHL3 residues Tyr449 and Phe402, where it can hydrogen bond to KLHL3 Ser432. Finally, the backbone carbonyls of WNK3 Cys538 and Glu539 form conserved hydrogen bond interactions between KLHL3 side chains Arg360 and Arg339, respectively.

**Figure 5.**
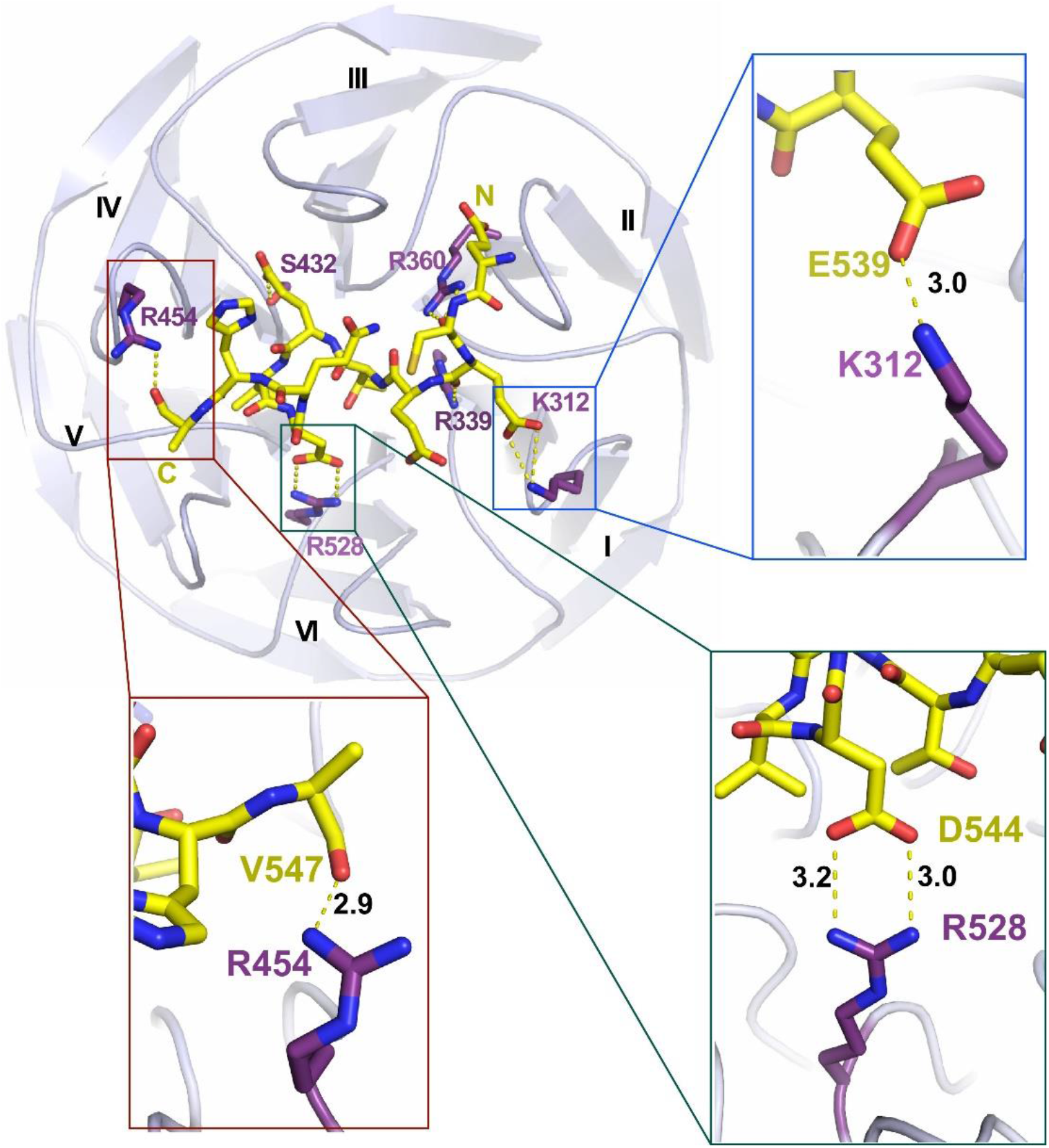
Hydrogen bond interactions in the KLHL3-WNK3 interface. Overview of the polar contacts in the complex interface. KLHL3 residues are labelled in purple and WNK3 peptide is shown in yellow. Hydrogen bonds are demonstrated using dashed lines. Inset panels show selected contacts. Hydrogen bond distances (Å) are indicated.

### Newly observed interactions in the WNK3 binding interface

The new structure reveals several features of the WNK family degron interaction that were not previously defined. First, the full 11-mer degron motif can be traced in the WNK3 structure, whereas the two C-terminal residues were not observed in the WNK4 complex. Surprisingly, truncation of either of these residues was not tolerated in WNK4 showing their importance for the KLHL3 interaction [9]. The newly defined C-terminal residues, WNK3 His546 and Val547, appear flexible to pack on the surface of the kelch domain forming van der Waals and electrostatic interactions with KLHL3 Tyr449 and Arg454, including a hydrogen bond between the backbone carbonyl of WNK3 Val547 (chain C) and KLHL3 Tyr449 (Figure 5). They also contribute to the peptide’s helical turn conformation through the backbone interactions of His546. Second, the new structure reveals a salt bridge between the conserved WNK3 residue Glu539 and KLHL3 Lys312 (Figure 5). The side chain atoms of this glutamate residue appeared disordered in the WNK4 structure, possibly resulting from its solvent exposed position and the additional flexibility of the associated lysine.

### Steric constraints would disfavour phosphothreonine in the WNK3 interface

WNK3 Thr541 adopts a central buried position in the complex interface and would appear to offer a unique potential phosphorylation site for regulation among the WNK family degron motifs (Figures 6A and 6B). The threonine side chain forms van der Waals interactions with several KLHL3 residues, including Arg339 and Tyr577 (Figures 6C and 6D). While too distant from KLHL3 Ser433 and Arg339 to form direct hydrogen bonds, there is also potential for these side chains to bind through water-mediated interactions that are not resolved at the 2.8 Å resolution of the structure.

**Figure 6.**
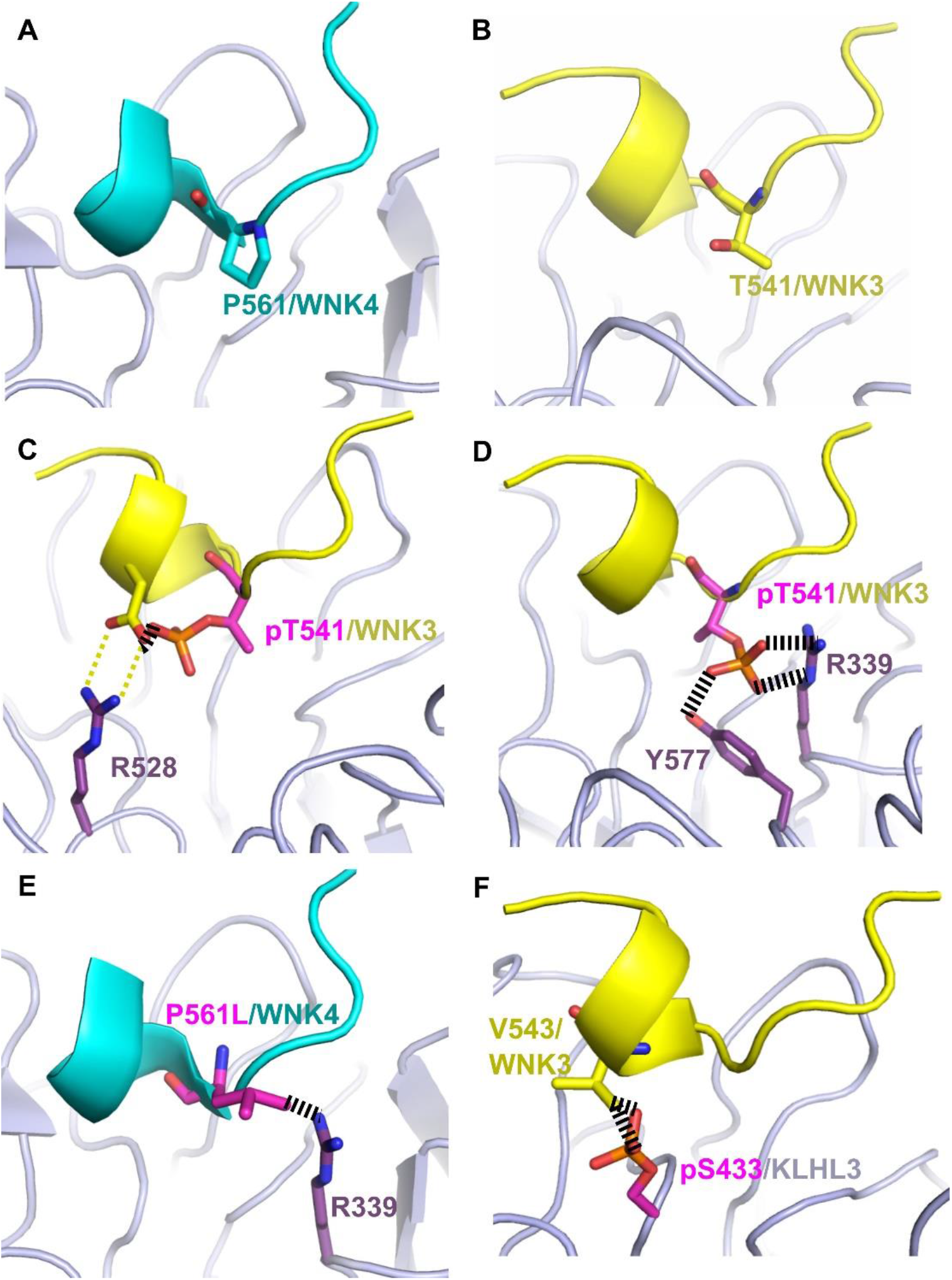
Steric constraints would disfavour phosphothreonine in the WNK3 interface. (**A**) Location of WNK4 Pro561 at the centre of the KLHL3-WNK4 binding interface (PDB 4CH9). KLHL3 is shown in light purple and WNK4 is shown in light blue. (**B**) Similar position of WNK3 Thr541 in the structure of the KLHL3-WNK3 complex (PDB 5NKP). WNK3 is shown in yellow. (**C**) The modelling of a phosphothreonine in the crystallized conformation of WNK3 Thr541 reveals a severe steric clash with WNK3 Asp544 that would break the critical salt bridge (yellow dashed lines) between this aspartate and KLHL3 Arg528. (**D**) Alternative binding pose of WNK3 Thr541 suggested after an energy minimization procedure to explore other potential side chain conformations. In this alternative pose, the phosphate moiety of pThr541 packs between KLHL3 Arg339 and Tyr577 in a sterically crowded environment that would bury the negative charge unfavourably without hydrogen bonding. (**E**) The modelling of a WNK4 PHAII mutation P561L reveals a steric clash (broken lines) with KLHL3 Arg339 that would disrupt the KLHL3-WNK4 interaction. **(F)**The modelling of Akt and PKA-dependent phosphorylation of KLHL3 Ser433 shows a severe steric clash with WNK3 Val543 that impairs KLHL3-WNK3 binding. The modelled residues in panels C-F are shown in pink.

To understand how post-translational modification on Thr541 might affect WNK3 binding to KLHL3, we modelled a phosphate moiety onto the threonine side chain using the ICM-Pro software package (Molsoft)[19]. When modelled in its crystallized conformation a severe steric clash was observed between pThr541 and WNK3 Asp544 that would break the critical salt bridge between this aspartate and KLHL3 R528 (Figure 6C). We therefore performed an energy minimization step to explore other potential side chain conformations. The resulting alternative conformation positioned the phosphate moiety of pThr541 between Arg339 and Tyr577 in a sterically crowded environment that would still bury the negative charge unfavourably without satisfying the required hydrogen bonding (Figure 6D). Thus, both the fluorescence polarisation assay and structural analysis indicate that Thr541 phosphorylation would disrupt the WNK3-KLHL3 interaction and therefore serve as a potential regulatory mechanism.

The WNK3 degron motif most closely matches the preferred substrate sites of acidophilic kinases such as casein kinase II (CK2) [20]. However, to date there have been no reports of phosphorylation on WNK Thr541 either in the literature or in public databases. The majority of proteomic analyses use trypsin digestion to restrict cleavage sites to basic arginine and lysine positions, which simplifies database searches for peptide matches. We observed that WNK3 Thr541 lies in a particularly poor sequence region for this approach as trypsin digestion would yield a peptide 36 amino acids in length due to the paucity of surrounding basic residues. We reasoned therefore that a potential phosphorylation at this site may have been missed previously and decided to perform a new proteomic search using elastase cleavage which targets the more common positions of small hydrophobic residues [21]. For these experiments, we transfected HEK293T cells with Flag-tagged WNK3 plasmid and after 36 hours treated the cells for 30 minutes with either isotonic or hypotonic buffer to allow for phosphorylation under different stimuli. Samples enriched for WNK3 were then prepared by anti-Flag immunoprecipitation, cleaved and analysed by mass spectrometry. Using this approach we obtained near complete coverage of the WNK3 sequence including the degron motif (Figures S1 and S2). Reassuringly, the experiment identified the known phosphorylation site within the kinase activation segment at Thr308. However, no phosphorylation was detected at the degron site residue Thr541 (Figures S1 and S2).

Overall, these results indicate that the four amino acid substitutions within the WNK3 acidic degron motif compared to WNK4 are well tolerated with no significant changes in the bound peptide conformation or binding affinity. We identify WNK3 Thr541 as a potential site for regulation, but further proteomic screening will be needed to demonstrate whether this site is phosphorylated under certain stimuli or in specific cell types.

## Discussion

Relatively few of the ~600 human E3 ligases have been structurally characterised in complex with a bound substrate and only a small subset of these have had their structures solved with multiple partners to investigate potential differences in binding modes and regulation. To date, the only example within the BTB-Kelch family has been the redox sensor KEAP1 (KLHL19), which has been crystallised in complex with multiple partners, including the substrate Nrf2 [22–25] and the substrate competitors prothymosin alpha [26] and p62 (Sequestosome-1) [27, 28]. These studies have defined notable substrate differences. For example, the divergent ‘ETGE’ and ‘DLG’ degron sites in Nrf2 have shown distinct binding affinities and bound peptide conformations [22]. Comparable KEAP1 structures have also been solved in complex with the related ‘STGE’ motif of p62 in both its serine phosphorylated [27] and non-phosphorylated forms [28]. In these structures, the phospho-serine provided p62 with enhanced binding affinity by mimicking the first glutamate of the Nrf2 ETGE motif [27].

Multiple substrates have been similarly identified for the KLHL3 E3 ligase, including the four members of the WNK kinase family, WNK1-4 [7–9]. WNK kinases display overlapping, but non-redundant functions reflecting with their different tissue expression profiles and regulation [1, 4]. For example, loss of WNK3 may reduce ischemia-associated brain damage [29], whereas mutations in WNK1 and WNK4 disrupt renal salt uptake to induce pseudohypoaldosteronism type II (PHAII) [6]. Here, we investigated the variant degron motif of WNK3. Our fluorescence polarisation data confirmed that WNK3 binds to KLHL3 with comparable affinity to other WNK isoforms despite carrying four amino acid substitutions within its degron. These data also revealed that the binding was dependent on WNK3 Thr541. A mutation at the equivalent position in WNK4, P561L, causes PHAII showing the importance of this position for WNK kinase regulation [30] (Figure 6B).

The four substitutions in WNK3 remove a PXXP sequence motif that is conserved in the degrons of WNK1, WNK2 and WNK4 raising the question of whether the complex between WNK3 and KLHL3 may adopt a distinct binding mode. However, the new structure of the KLHL3-WNK3 complex reveals that the WNK3 peptide binds across the Kelch domain surface with a binding pose that is conserved with the equivalent WNK4 complex. In fact, the main difference appears to be a subtle shift at the binding interface to accommodate an alanine to valine substitution in WNK3, as well as WNK3 Thr541. The new structure also reveals several features of the degron interaction that were not previously defined. These include the salt bridge formed by the conserved WNK3 residue Glu539 and the additional packing interactions of the two C-terminal degron positions (WNK3 His546 and Val547) that contribute to the peptide’s helical turn conformation.

Akt and PKA-dependent phosphorylation of KLHL3 Ser433 can impair the recruitment and ubiquitin-dependent degradation of WNK kinases [31, 32]. Our structure reveals that this KLHL3 residue is positioned in close proximity to the bound WNK3 residues Val543 (3.85 Å) and Thr541 (4.9 Å), which would not tolerate the phospho-serine moiety due to a severe steric clash (Figure 6F). We hypothesised that a similar regulatory mechanism may exist in the WNK family through the phosphorylation of WNK3 Thr541. Indeed, structural modelling and our fluorescence polarisation binding assay both indicated that this phosphorylation would disrupt the KLHL3-WNK3 interaction. Under hypotonic conditions, WNK3 mediates the inhibitory phosphorylation of KCC3 T991 and T1048 to restrict cell swelling in the brain [29, 33]. We speculated whether such hypotonic conditions might promote the phosphorylation of WNK3 Thr541 to modulate WNK3 stability. Unfortunately, our proteomic studies were unable to detect evidence of pThr541 in HEK293T cells under either hypotonic or isotonic treatment. However, we cannot exclude that such phosphorylation occurs under other stimuli or in other specific cell types.

Overall, the current work extends our understanding of the molecular mechanisms that control WNK kinase ubiquitination and degradation. It shows that the four WNK family kinases share a conserved mechanism of binding to KLHL3 through an 11-mer acidic degron motif and highlights WNK3 Thr541 as a potential regulatory site for the E3 ligase interaction.

## FUNDING

The SGC is a registered charity (number 1097737) that receives funds from AbbVie, Bayer Pharma AG, Boehringer Ingelheim, Canada Foundation for Innovation, Eshelman Institute for Innovation, Genome Canada, Innovative Medicines Initiative (EU/EFPIA) [ULTRA-DD grant no. 115766], Janssen, Merck KGaA Darmstadt Germany, MSD, Novartis Pharma AG, Ontario Ministry of Economic Development and Innovation, Pfizer, São Paulo Research Foundation-FAPESP, Takeda, and Wellcome [106169/ZZ14/Z]. D.R.A is supported by the Medical Research Council [grant number MC_UU_12016/2] and the pharmaceutical companies supporting the Division of Signal Transduction Therapy Unit (Boehringer-Ingelheim, GlaxoSmithKline, Merck KGaA -to D.R.A.).

## ACKNOWLEDGMENTS

The authors would like to thank Diamond Light Source for beamtime (proposal mx10619), as well as the staff of beamline I03 for assistance with crystal testing and data collection. Mass spectrometry analysis was performed at the Discovery Proteomics Facility (headed by Roman Fischer) which is part of the TDI MS Laboratory (led by Benedikt Kessler).

## Author Contributions

A.N.B. and D.R.A. designed the research. J.Z. performed the binding assays. Z.C. purified and crystallised the KLHL3-WNK3 complex and refined its structure with assistance from F.J.S. Z.C. performed tissue culture and sample preparation for phosphorylation mapping with assistance from V.D.A. LC-MS/MS was performed by R.H. and R.F. The initial manuscript draft was prepared by A.N.B. and Z.C. and completed with the help of all authors.

## Competing Interests

The Authors declare that there are no competing interests associated with the manuscript.

## Abbreviations

BTB: Bric-a-brac, Tramtrack, and Broad complex
CCT: Conserved C-terminal
CRL3: Cullin3-dependent RING E3 ligase
DMEM: Dulbecco’s modified Eagle’s medium
DTT: Dithiothreitol
FBS: fetal bovine serum
GST: Glutathione S-transferase
HCD: Higher-energy collisional dissociation
HEK293T: human embryonic kidney 293 cell line expressing the SV40 large T antigen
KCC: K+/Cl− co-transporters
IPTG: Isopropyl β-D-1-thiogalactopyranoside
KLHL3: Kelch-like protein 3
LC/MS: Liquid chromatography-mass spectrometry
N[K]CC: Na+/Cl− ion co-transporters
NKCC2: Na+/K+/2Cl− co-transporter 2
OSR1: oxidative stress-responsive kinase 1
PDB: Protein Databank
PHAII: pseudohypoaldosteronism type II
RING: really interesting new gene
SPAK: SPS1-related proline/alanine-rich kinase
TCEP: tris(2-carboxyethyl)phosphine
TEV: tobacco etch virus
WNK: with no lysine (K) kinase

## Notes

### Competing Interest Statement

The authors have declared no competing interest.

