## Supplementary Figures S1-S2 for "Sequence and structural variations determining the recruitment of WNK kinases to the KLHL3 E3 ligase"

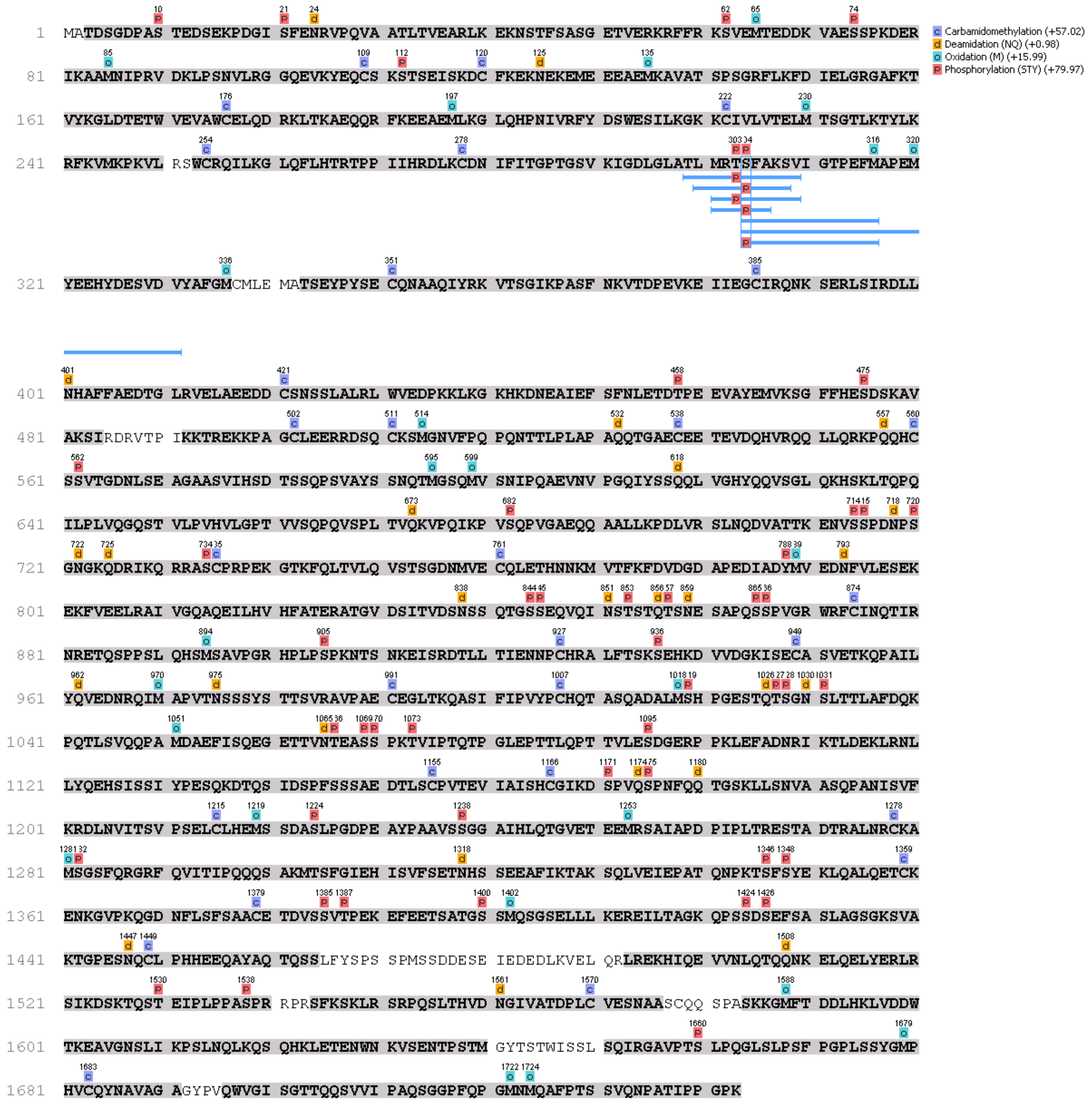


**Figure S1. Overview of WNK3 peptide hits from hypotonic treatment.** Residues recovered in elastase MS/MS are shaded in grey. Modifications are labelled above residues. Peptide hits in the kinase activation segment are illustrated.

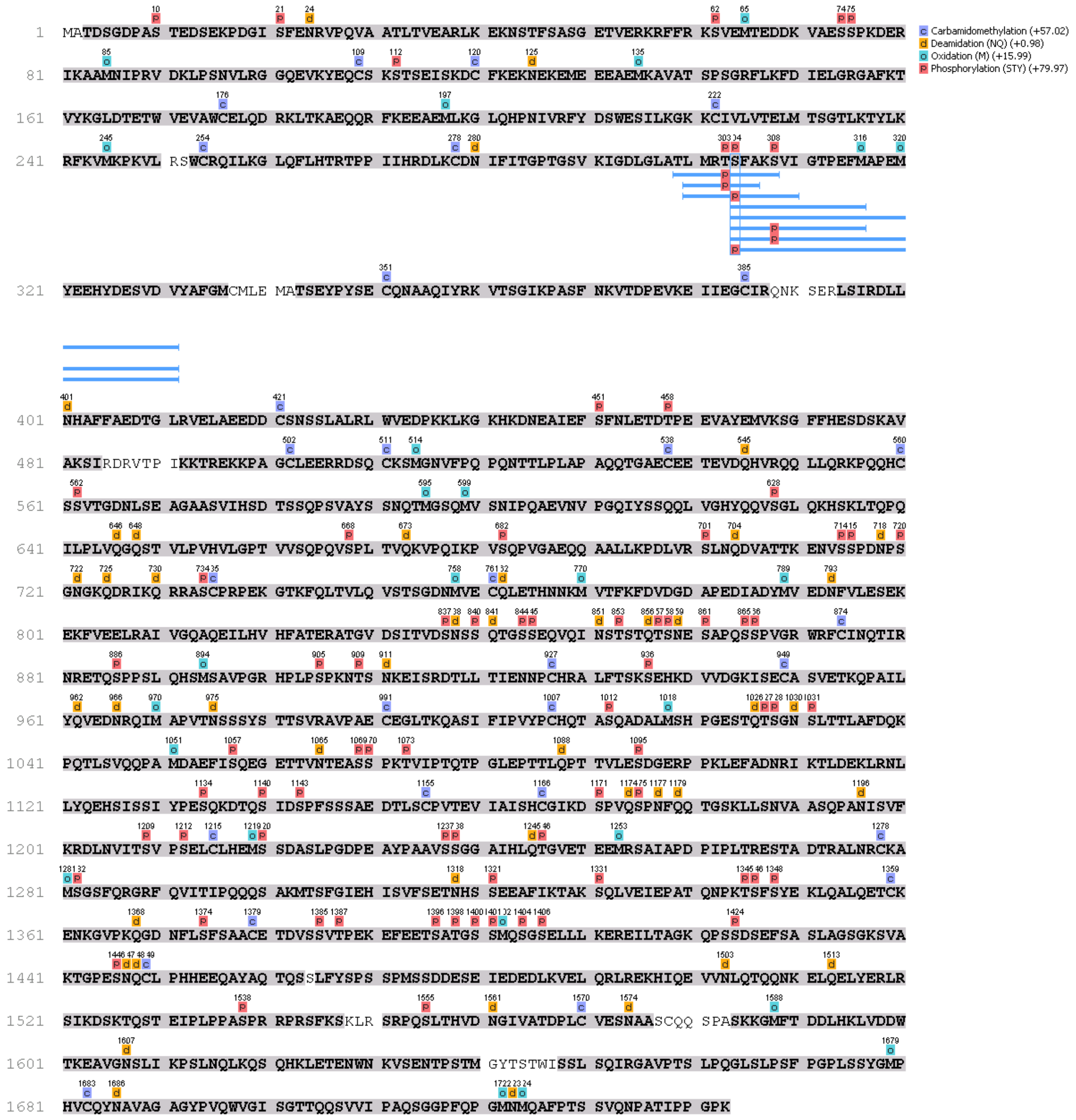

**Figure S2. Overview of WNK3 peptide hits from isotonic treatment.** Residues recovered in elastase MS/MS are shaded in grey. Modifications are labelled above residues. Peptide hits in the kinase activation segment are illustrated.
